# Strong and consistent associations of waterbird community composition with HPAI H5 occurrence in European wild birds

**DOI:** 10.1101/2022.04.11.487853

**Authors:** Zheng Y.X. Huang, Shenglai Yin, Yong Zhang, Willem F. de Boer, Taej Mundkur, Jean Artois, Francisca Velckers, Huaiyu Tian, John Y. Takekawa, Yali Si, Guan-Zhu Han, Huimin Zhang, Yuyang Chen, Hongliang Chai, Chi Xu

## Abstract

Since 2014, highly pathogenic avian influenza (HPAI) H5 viruses of clade 2.3.4.4 have been dominating the outbreaks across Europe, causing massive deaths among poultry and wild birds. However, the factors shaping its broad-scale outbreak patterns remain unclear. With extensive waterbird survey datasets of about 7,000 sites across Europe, we here demonstrated that H5N8 occurrence in wild birds in the 2016/17 and 2020/21 epidemics as well as H5N1 occurrence in 2005/06 epidemic were strongly associated with very similar waterbird community attributes, pointing to the possibility of similar interspecific transmission processes between different epidemics. A simple extrapolation of the model constructed from the 2016/17 epidemic can well predict the H5N8 pattern in wild birds in 2020/21 epidemic. We also found a dilution effect of phylogenetic diversity that was always negatively correlated with H5 occurrence in wild birds. In contrast, H5N8 occurrence in poultry was subject to different risk factors between the two epidemics. In general, waterbird community composition play a much more important role in determining the spatial pattern of H5N8 in wild birds than in poultry. Our work contributes to reveal the factors driving H5N8 patterns, and highlights the value of waterbird community factors in future HPAI surveillance and prediction.

## Introduction

Emerging infectious diseases are one of the most critical challenges to humanities. The pandemic of COVID-19 has attracted the world’s great attention and medical resources, but shadows over many others in recent years. Over the last two decades, outbreaks of highly pathogenic avian influenza (HPAI), mostly caused by H5 viruses, have occurred frequently in European countries (Verhagen et al. 2021). Europe has experienced several severe outbreaks (Alarcon et al. 2018, Verhagen et al. 2021), including those in 2005/06 H5N1 (clade 2.2), 2014/15 H5N8 (clade 2.3.4.4a), 2016/17 H5N8 (clade 2.3.4.4b) and 2020/21 H5N8 (clade 2.3.4.4b) epidemics. These have caused great mortality in wild birds and poultry (Lycett et al. 2019, Verhagen et al. 2021), and posed threats to public health (Pyankova et al. 2021, Shi and Gao 2021). Previous studies have suggested that broad-scale spatiotemporal patterns of H5N1 occurrence appear largely driven by agro-climatic factors such as human population density, poultry density, cropping intensity, distance to waterbodies, temperature and precipitation (Gilbert and Pfeiffer 2012). However, it is unclear if these factors also can drive H5N8 (and other subtypes) distributions as only very few studies have analyzed H5N8 distributions (but see (Kim et al. 2018)). Addressing this knowledge gap is particularly important for the prediction and prevention of future HPAI outbreaks.

Recent work increasingly highlights that local-scale biotic interactions can produce important cross-scale effects to drive broad-scale ecological patterns (Teng et al. 2020). Such cross-scale effect may also hinge on the formation of continental patterns of HPAI. For example, seasonal migration of waterbirds has been suggested to play a critical role in the long-distance spread of H5N1 (Tian et al. 2015, Xu et al. 2016) and H5N8 (Global Consortium for H5N8 and Related Influenza Viruses 2016), implying that waterbirds may act as a key nexus of local-scale HPAI outbreaks and broad-scale epidemic patterns. However, there is relatively little evidence to support this theory. For example, a study explored the risk factors for H5N1 occurrence during the 2005/06 epidemic in Europe, and suggested that local-scale waterbird community composition could effectively explain the continental pattern of H5N1 occurrence in wild birds (Huang et al. 2019b). It is highly uncertain if local waterbird community factors played a similar role in other HPAI subtypes, as H5N8 epidemic appeared to have clearly different features than the H5N1 in terms of the epidemic curve as well as duration and intensity (Alarcon et al. 2018). Important gaps remain for identifying risk factors of H5N8 outbreaks, especially when it comes to the role of waterbirds as a critical component of transmission pathways.

In this paper, we investigate the spatial patterns of H5N8 outbreaks during the 2016/17 and 2020/21 epidemics in Europe. We applied waterbird census data from about 7,000 sites across Europe to 1) explore and compare the risk factors associated with H5N8 occurrence in wild birds and poultry, and between the 2016/17 and 2020/21 epidemics; and 2) determine the importance of waterbird community factors in predicting H5N8 spatial patterns.

## Results

The H5N8 data collected from FAO (see Methods) included 1,630 outbreak cases from the 2016/17 epidemic with 778 cases in wild birds and 852 in poultry (Figure 1A), and included 1,581 outbreak cases reported from the 2020/21 epidemic with 743 cases in wild birds and 838 cases in poultry (Figure 1B).

**Figure 1:**
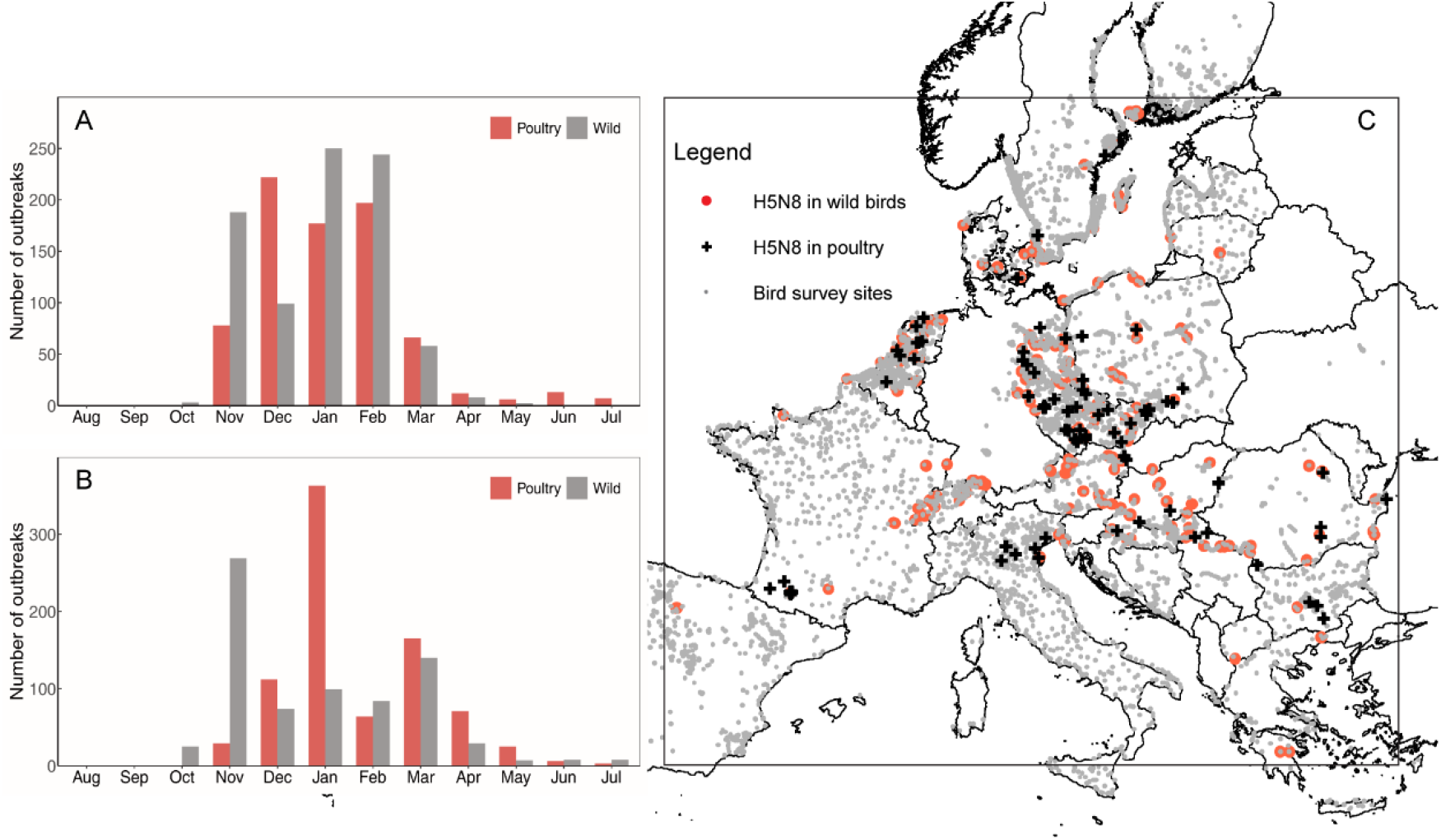
Temporal distribution of H5N8 outbreak cases for poultry and wild birds in Europe during the 2016/17 epidemic (A) and 2020/21 epidemic (B) and spatial distribution of H5N8 outbreaks and bird survey sites during the 2016/17 epidemic (C).

### Risk factors for H5N8 occurrence

We conducted univariate analyses to explore risk factors for H5N8 occurrence in the 2016/17 and 2020/21 epidemics. The factors associated with H5N8 occurrence in wild birds were similar between the two epidemics (Figure 2A). H5N8 in wild birds was negatively correlated with climatic variables (i.e., temperature, precipitation) and community phylogenetic diversity indices, while positively correlated with chicken density, the total length and area of waterbodies in the buffer zone, and several abundance-related variables including the abundance of Mallard (*Anas platyrhynchos*), Eurasian Wigeon (*A. penelope*), Tufted Duck (*Aythya fuligula*), Mute Swan (*Cygnus olor*) and all waterbirds. The community risk factors for H5N8 in wild birds were also similar to those for H5N1 occurrence in wild birds in the 2005/06 epidemic, except for the abundance of Black-headed Gull (*Larus ridibundus*) that showed a positive relationship with H5N1 in wild birds (*Supplementary file 1a*). In contrast, the risk factors for H5N8 occurrence in poultry were different between the 2016/17 and 2020/21 epidemics (Figure 2B), except for climatic variables and chicken density, which respectively had negative and positive correlations in both epidemics. Duck density and several abundance-related variables (i.e., the abundance of bridge species, Mallard and Eurasian Wigeon) were positively correlated with outbreak probability in the 2016/17 epidemic, while none of them showed a significant relationship in the 2020/21 epidemic.

**Figure 2:**
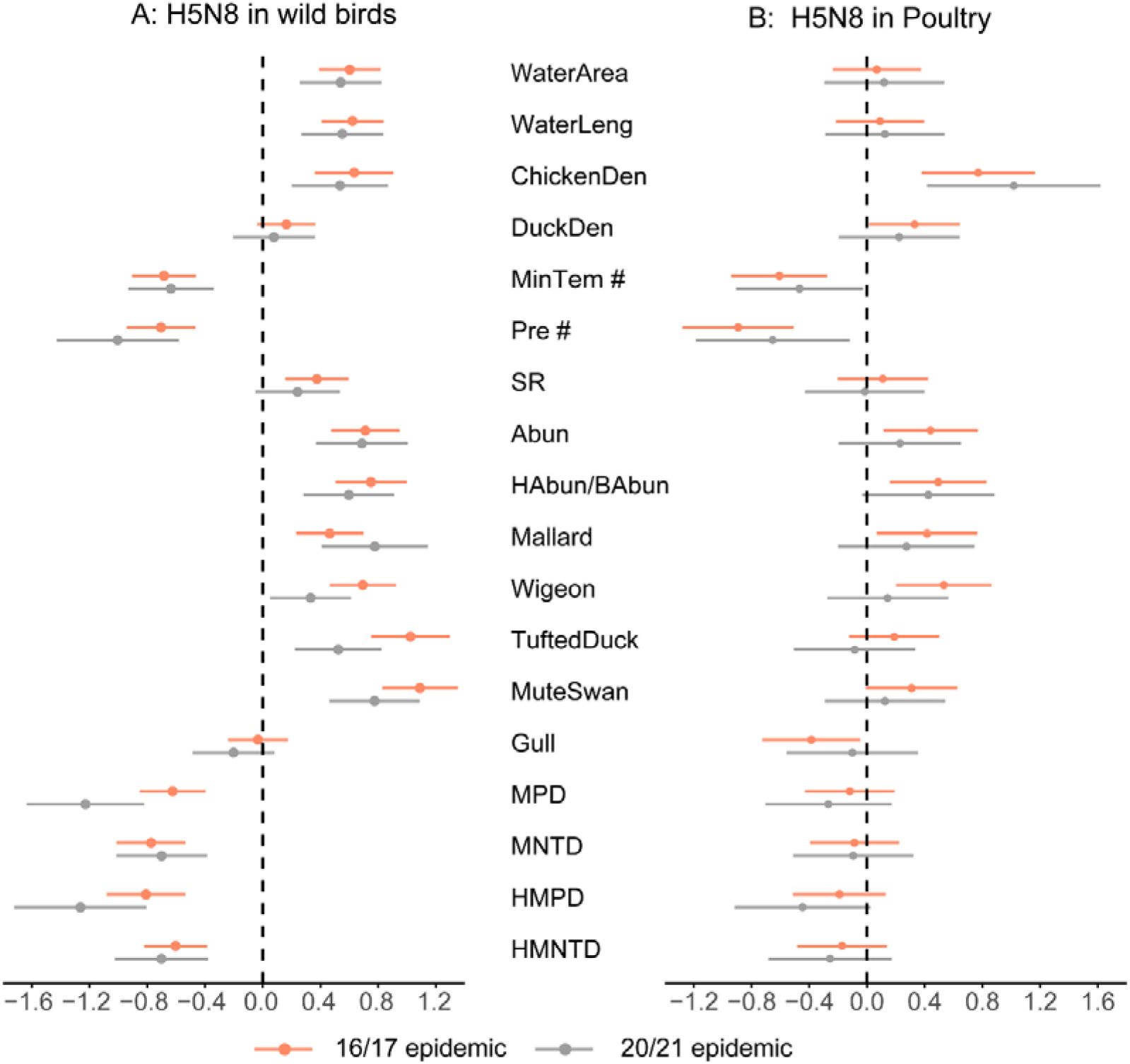
Results (regression coefficient, b ± 95% CI) of univariate regression analyses for H5N8 outbreaks in wild birds (A) and in poultry (B) in the 2016/17 and 2020/21 epidemics where variables with CIs overlapping 0.0 are uncorrelated. Agro-climatic variables included the total area (WaterArea) or length (WaterLeng) of waterbody, chicken density (ChickenDen), domestic duck density (DuckDen), the mean monthly minimum temperature (MinTem) and the mean monthly precipitation (Pre). Community variables included waterbird species richness (SR), the abundances of total waterbirds (Abun), high-risk or bridge species (HAbun/BAbun), mallard (Mallard), Eurasian Wigeon (Wigeon), Tufted Duck (TuftedDuck), Mute Swan (MuteSwan) and Black-headed Gull (Gull), the standardized mean pairwise distance (MPD), standardized mean distances of nearest neighbours (MNTD) and these two metrics based on high-risk species (HMPD and HMNTD). # Only the temperature and precipitation variable with the smallest p-values were shown in the figure

### Relative importance of community predictors in explaining H5N8 patterns

We then constructed variance partitioning analyses to compare relative importance of community versus agro-climatic predictors in determining the spatial patterns of H5N8 occurrence. For H5N8 outbreaks in wild birds, the variation solely explained by community predictors were, respectively, 16.5% and 17.8% for the 2016/17 and 2020/21 epidemics, whereas those explained by agro-climatic predictors were 15.5% and 12.4% (Figure 3). For the H5N8 outbreaks in poultry in the 2016/17 epidemic, the variation solely explained by community predictors and agro-climatic predictors were, respectively, 8.2% and 33.6% (Figure 3). We did not conduct variance partitioning analyses for H5N8 outbreaks in poultry in the 2020/21 epidemic, because none of the community factors showed a significant relationship in univariate analyses (Figure 2).

**Figure 3:**
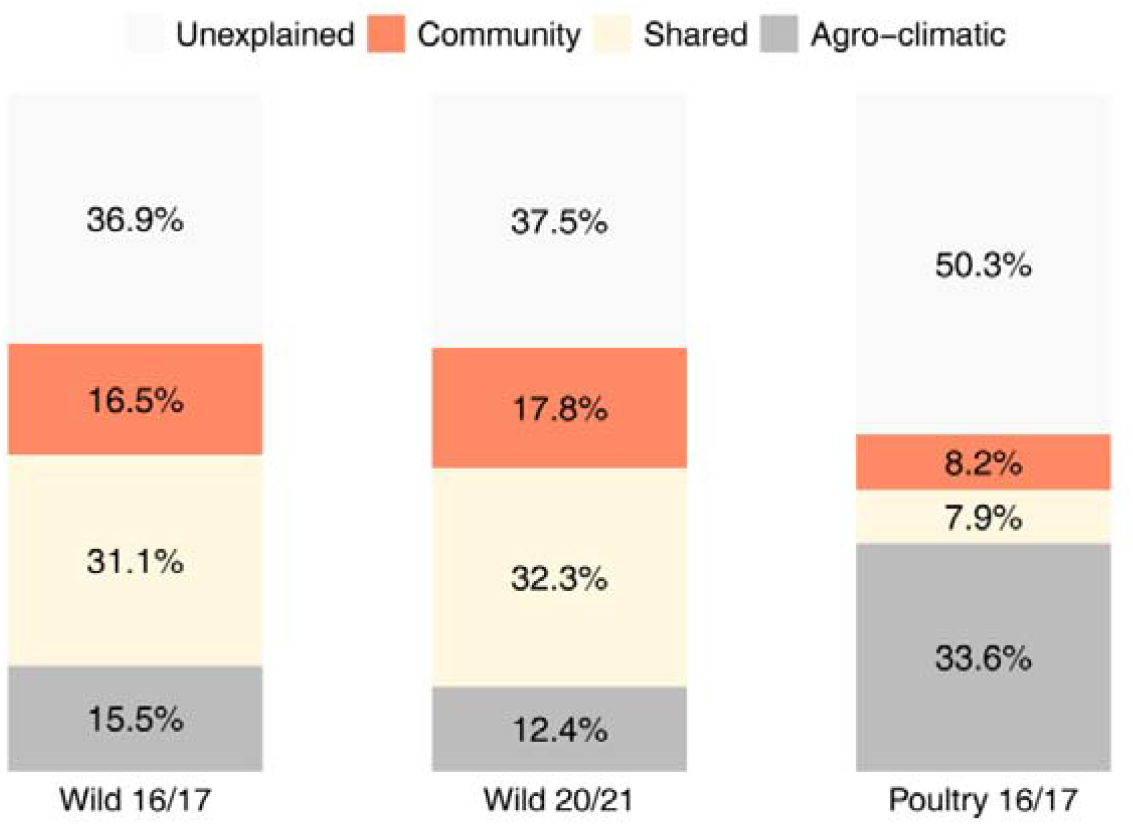
Variance partitioning analyses, illustrating the percentages of variance explained by different groups of predictors for H5N8 occurrence in wild birds in the 2016/17 and 2020/21 epidemics and H5N8 in poultry in the 2016/17 epidemic.

### Predictive powers of the models based on previous data for recent H5N8 patterns

We finally constructed multiple regression models for H5N8 in the 2016/17 epidemic (training dataset) and determined their predictive powers for the 2020/21 epidemic (test dataset). The final model for H5N8 in wild birds had an AUC value of 0.913 for the training dataset and 0.819 for the test dataset. This final model included seven predictors (Figure 4), with positive correlations for total length of waterbodies, chicken density, and abundances of Tufted Duck and Mute Swan, and negative correlations for mean precipitation in the winter, temperature in January, and pairwise phylogenetic distance of high-risk species. The final model for H5N8 in poultry had an AUC value of 0.848 for the training dataset and 0.598 for the test dataset, and it included four predictors, namely duck density and the abundance of Eurasian Wigeon with positive correlations, and mean precipitation in February and minimum temperature in January with negative correlations. In addition, the relationships identified in the final models were consistent in sensitivity analyses (*see Materials and methods*) where we changed the thresholds for identifying pseudo-absences (*Supplementary file 1b and 1c*) or removed the poor matches in presences (*Supplementary file 1d*).

**Figure 4:**
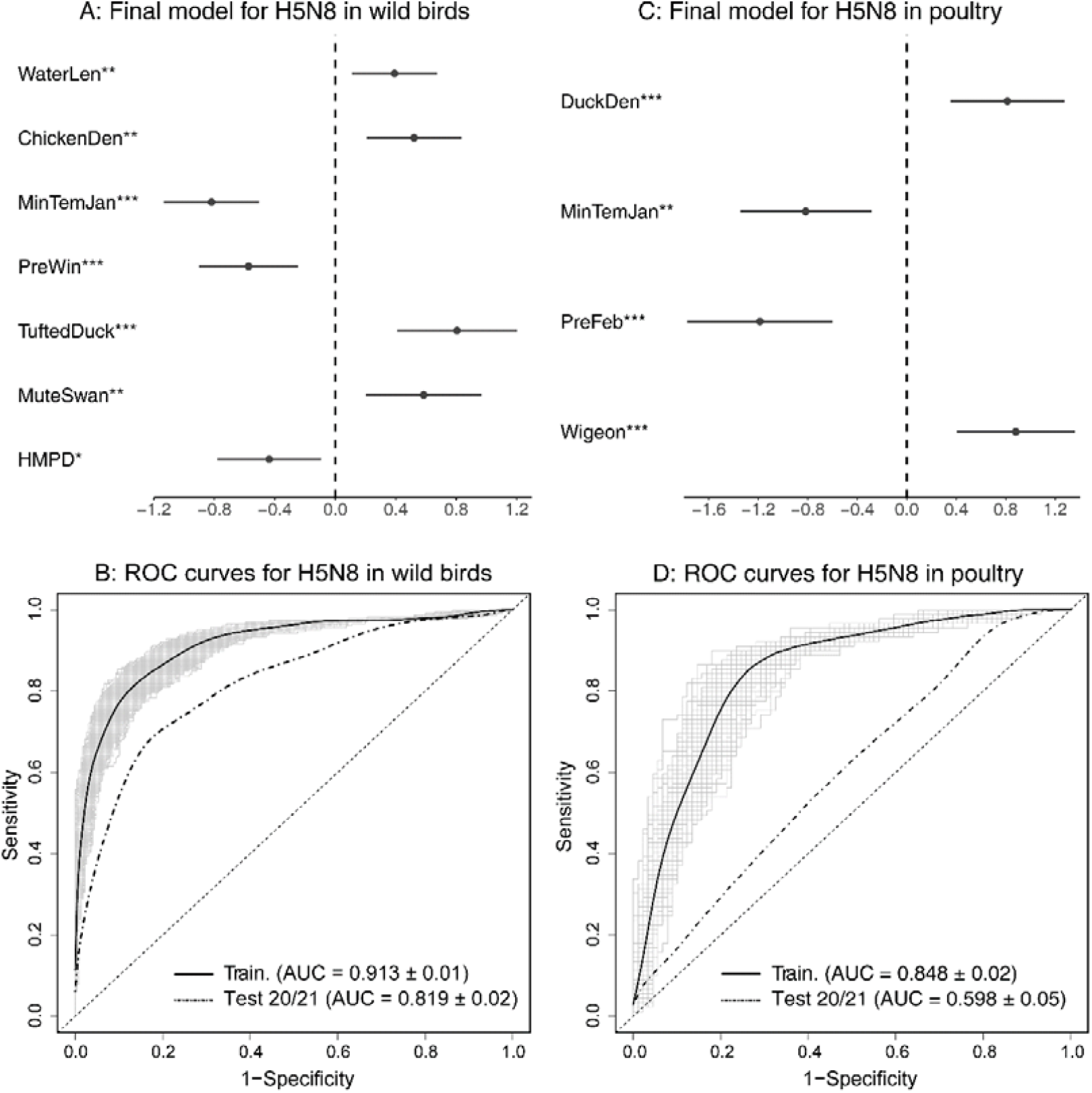
Results of the final model predicting the presence/absence of HPAI H5N8 in wild birds (A and B) and poultry (C and D). A and C: regression coefficients (b ± 95%CI); B and D: mean ROC curves (with 1000 ROC curves for training data) and AUC values. Variables included the total length (WaterLeng) of waterbodies, chicken density (ChickenDen), domestic duck density (DuckDen), mean minimum temperature in January (MinTemJan) and mean precipitation in February (PreFeb), abundances of Eurasian Wigeon (Wigeon), Tufted Duck (TuftedDuck), Mute Swan (MuteSwan) and Black-headed Gull (Gull), and standardized mean pairwise distance of high-risk species (HMPD).

## Discussion

Our analyses demonstrate that waterbird community composition was strongly associated with the spatial patterns of HPAI H5N8 occurrence in wild birds. Importantly, the risk factors and their effects were highly consistent between the two epidemics. A simple extrapolation of the model constructed from the 2016/17 epidemic can predict the 2020/21 epidemic pattern well. In contrast, the factors driving H5N8 patterns in poultry were different than in wild birds, and inconsistent between the two epidemics.

In both epidemics, the community predictors were as important as agro-climatic predictors in explaining the spatial pattern of H5N8 occurrence in wild birds. The abundances of Tufted Duck, Mute Swan and two *Anas* species (Mallard and Eurasian Wigeon) were positively correlated with H5N8 occurrence in wild birds. A previous study identified Tufted Duck and Mute Swan as the species most frequently infected with H5N8 during the 2016/17 epidemic in Europe, and considered them as local amplifiers of H5N8 virus because they rarely fly between foraging lands and roosting sites on their wintering grounds (Alarcon et al. 2018). Another study analyzing the deaths of wild birds in the Netherlands during the 2016/17 epidemic also found that Tufted Duck and Eurasian Wigeon were the most recorded species that died (Kleyheeg et al. 2017), suggesting that they are particularly susceptible to the H5N8 clade 2.3.4.4b.

Apart from these factors related to particular species, community level factors correlated with H5N8 occurrence in both epidemics in a consistent way. Host phylogenetic diversity showed a negative relationship, suggesting that communities with closely related host species exhibit a relatively higher disease risk. Previous studies suggested closely related species might have similar immunological responses and life-history traits, which increase the probability of interspecific transmission and infection (Liu et al. 2016, Wang et al. 2019b). In addition, closely related species may aggregate at similar habitats due to shared physiological or behavioral requirements (Fountain-Jones et al. 2018), leading to a high contact rate among hosts and virus and increasing disease risk. Our results also provide empirical support for the potential existence of a dilution effect of phylogenetic diversity that only very few studies has considered (bet see (Huang et al. 2019a, Wang et al. 2019a)).

Strikingly, the abovementioned risk factors for H5N8 in wild birds were also responsible for the 2005/06 H5N1 epidemic. Although a previous study concluded that the 2016/17 H5N8 epidemic was different from the 2005/06 H5N1 epidemic in several aspects (e.g., epidemic curve, outbreak duration, and poultry illness and death rate)(Alarcon et al. 2018), the outbreak distributions among different waterbird species (i.e., the percentage of outbreak events in different species) were similar between the two epidemics (Table 2 in (Alarcon et al. 2018)). Our finding provide an important suggestion that the interspecific transmission processes among wild bird species could be similar between the three epidemics, even different H5 subtypes were involved.

Our analyses demonstrated that the risk factors for H5N8 occurrence in poultry were different than those in wild birds and were not consistent between the 2016/17 and 2020/21 epidemics. In line with previous studies (Kim et al. 2018, Napp et al. 2018, Briand et al. 2021, Gierak and Śmietanka 2021), we found a positive relationship of domestic duck density in the 2016/17 epidemic. We further found that in 63 sites with reported chicken outbreaks in the 2016/17 epidemic, 26 sites experienced domestic duck outbreaks within two weeks prior in the surrounding area (< 30 km), while none of the 410 duck outbreak sites experienced any prior outbreaks in other domestic species. These results reinforce that domestic ducks could be a key component in the maintenance and spread of the virus to chicken farms (Yoo et al. 2021).

In addition, domestic ducks have been considered to be key intermediate hosts between wild birds, particularly *Anas* species, and poultry (Cappelle et al. 2014, Kwon et al. 2020). We here also found that the abundances of two *Anas* species (i.e., Mallard and Tufted Duck) had positive effects in the 2016/17 epidemic. This could be because the two species usually forage on grasslands and agricultural lands which are often relatively close to farms (Kleyheeg et al. 2017, Belkhiria et al. 2018). In fact, a recent study also found that in the Netherlands the density of Eurasian Wigeon was considerably higher around farms infected by H5N8 compared to non-infected farms in the 2016/17 epidemic (Velkers et al. 2021). However, neither domestic duck density nor the abundance of *Anas* species showed a significant effect in the 2020/21 epidemic. In fact, there were much fewer cases reported in domestic ducks in the 2020/21 epidemic (13 cases), compared to the 2016/17 epidemic (410 cases), but the reasons for these differences between the two epidemics require further investigation. Local/regional analyses incorporating detailed socioeconomic data such as the wide range of poultry production systems in Europe should also be considered.

The databases from FAO probably underestimated the number of outbreaks, particularly wild bird outbreaks, due to the applied passive surveillance system (European Food Safety Authority et al. 2021), though these datasets have been widely used in studies mapping HPAI risk (Si et al. 2010, Martin et al. 2011, Si et al. 2013, Dhingra et al. 2016). In addition, the spatial mismatch between disease outbreak sites and waterbird survey sites may potentially influence our results. However, we considered that our results were robust, illustrated by similar results generated by the models using subsets of data, i.e. excluding the presences with poor matches (Table S8). Finally, due to the lack of more recent waterbird census data, we used waterbird data from 2015 to 2017 throughout the analyses, considering that the abundances of common species, such as Mallard, Eurasian Wigeon, Tufted Duck, and Mute Swan have remained relatively stable in recent years (iwc.wetlands.org). Future surveillance systems incorporating conditions of both diseases and ecological communities may effectively enhance our ability of understanding and predicting HPAI patterns.

Taken together, our work reveal the important and temporarily consistent role of waterbird community composition in driving the continental patterns of H5N8 occurrence in wild birds. Waterbird community composition played a larger role in H5N8 outbreaks in wild birds than that in poultry. Risk factors for H5N8 occurrence in poultry varied between the 2016/17 and 2020/21 epidemics. Our results imply that models based on previous epidemics might be predictive for future patterns of HPAI in wild birds but are less useful in predicting patterns in poultry. Our work contributes to understand the different H5N8 patterns in wild birds and poultry, and highlights the value of waterbird community factors in future HPAI surveillance and prediction.

## Materials and methods

### Data collection

We collected data of H5N8 outbreaks, together with their coordinates, in mainland Europe (Russia excluded) during the 2016/17 epidemic (from August 2016 to July 2017 as a whole epidemiologic year) and the 2020/21 epidemic from the databanks of the Food and Agriculture Organization of the United Nations (FAO) and removed duplicate records. We only analyzed the outbreaks between November and March, when bird census data were collected. The outbreak cases in this period accounted for, respectively, 96.9% and 88.5% of the total cases for the 2016/17 and 2020/21 epidemics.

Data on waterbird (species and counts) during 2015-2017 at about 7000 surveillance sites in 26 European countries (*Supplementary file 1e*) were obtained from the International Waterbird Census programme (IWC) coordinated by Wetlands International. In the northern wintering season, usually in mid-January but partly also in December or February, a large group of volunteers carried out the survey using standard census methods produced by Wetlands International. Thus, the method used for survey is consistent across sites, and the survey time matches well with the period when the majority of outbreaks occurred. For some surveillance sites, the survey was not carried out in all three years. For sites with only data from a one-year survey, we used the survey data of that particular year to represent the waterbird community, while the maximum number for any waterbird species was used for the sites with more than one years’ data. We also excluded rare species—those occurring in < 10 sites or with counts < 20 birds. These approaches have been widely used in many studies on waterbird community analyses.

### Matching waterbird community data to H5N8 outbreak sites

As bird surveillance sites usually are not matched exactly with the H5N8 outbreak sites, we here applied a cross-selection procedure to solve this spatial mismatch and generate disease “presences” and “pseudo absences” for further analyses. To achieve this, we first searched all surveillance sites around each outbreak site within a 10-km radius, which is related to surveillance zones for HPAI outbreaks in EU countries (Pittman and Laddomada 2008), and then selected the nearest surveillance site within this radius. Then, the bird census data for this site were assigned to the corresponding outbreak site as a “presence” site. We then classified these presence sites, according to the distance between bird surveillance site and outbreak site, as either 1) excellent match with distance less than 1 km; 2) good match with distance of 1–3 km; 3) fair match with distance of 3–5 km or 4) poor match with distance of 5–10 km. For the presence sites with fair and poor matches, we only included those where the surveillance site and H5N8 outbreak site were located around the same lake or river (less than 1 km away from the boundary of the lake or river). Moreover, for H5N8 outbreaks in wild birds, the selected surveillance site had to be located in a similar habitat (grassland or crop field based on land cover maps from the European Space Agency CCI 300-m annual global land cover products) as the H5N8 outbreak site. We used all matched presences to construct models but also tested the robustness by using data excluding the poor matches.

We then assigned surveillance sites sufficiently distant (> 60 km, three times the diameter for HPAI surveillance zones) from any outbreak site as potential H5N8 absence sites. We tested the sensitivity of our results by changing this distance threshold to 40 km and 80 km. We assumed the absences could represent a site unsuitable for infection, i.e., a true absence. However, there were, for spreading pathogens, also false absences resulting from a lack of surveillance and/or reporting, or because the pathogen had not been introduced into the region (Phillips et al. 2009). To limit the effect of these false absences, we also set a maximum distance to the nearest outbreak site in the selection of absence locations. We set the maximum distance at 200 km, and used 400 km and 800 km to test the consistency of our results. In addition, we only included absence sites in countries where H5N8 outbreaks occurred, and, like previous studies (Si et al. 2010, Huang et al. 2019b), within the maximum geographic range of H5N8 outbreaks (Figure 1).

### Processing of the predictors

For each presence or absence site, we calculated a set of community predictors related to the waterbird community composition (*Supplementary file 1f*). We considered several abundance-related variables, including the total abundance of waterbirds in the site (Abun), the abundances of bridge species (BAbun) which were identified by Veem et al. (Veen et al. 2007) based on species’ habitat, behavior and ecology (*Supplementary file 1g*), as well as the abundance of high-risk species (HAbun). We classified high-risk species by combining the list from Veen et al. (Veen et al. 2007) and the list from Schreuder (Schreuder 2021) (*Supplementary file 1g*). The variables BAbun and HAbun were, respectively, used for the analyses of H5N8 outbreaks in poultry and in wild birds. We also calculated the abundances of several important species, including Mallard (*Anas platyrhynchos*), Eurasian Wigeon (*Anas penelope*), Tufted Duck (*Aythya fuligula*), Mute Swan (*Cygnus olor*) and Black-headed Gull (*Larus ridibundus*), which were particularly shown to be infected with H5N8 virus (Kleyheeg et al. 2017, Napp et al. 2018, van den Brand et al. 2018, Schreuder 2021). We also included several diversity-related variables, including the species richness of waterbirds in the site (SR), and four variables measuring community phylogenetic diversity, as host relatedness within a community can affect pathogen transmission and disease risk (Liu et al. 2016, Huang et al. 2019b, Wang et al. 2019b). For each site, we calculated the standardized mean pairwise distance (MPD), standardized mean distances of nearest neighbors (MNTD) based on a consensus tree generated by summarizing 2,000 phylogenetic trees with a “Hackett” backbone (Hackett et al. 2008) from BirdTree (Jetz et al. 2012). These two indices measures represent, respectively, the overall average phylogenetic distance among species, and the overall average distance between closely related species within the community (Jetz et al. 2012, Tucker et al. 2017, Li et al. 2019). In addition, we also calculated MPD and MNTD based on assemblages only containing high-risk species (HMPD and HMNTD).

In addition to community predictors, we considered a large set of agro-climatic and anthropogenic predictors (*Supplementary file 1f*) that have been investigated previously. We calculated the monthly mean precipitation (Pre) and mean daily minimum temperature (MinT) for each month from December to February based on a 5-km-radius buffer zone around each sampling site, and averaged these mean variables over three months to get the averaged winter values. We then calculated the mean chicken density (ChickDen) and mean duck density (DuckDen), which have been identified as risk factors for both H5N1 (Gilbert and Pfeiffer 2012) and H5N8 occurrence (Dhingra et al. 2016, Briand et al. 2021, Gierak and Śmietanka 2021). Additionally, we calculated the total length and total area of waterbodies (WaterLen and WaterArea) in the buffer zone. Following previous studies (Gilbert et al. 2008, Dhingra et al. 2016), we calculated the mean human population density (PopDen) and the mean road density (RoadDen) within the buffer to control for the reporting bias, as under-reporting would be more likely in remote areas (Phillips et al. 2009). All data were collected from published databanks (see Data Availability) and were processed in ArcGIS v10.2.

### Statistical analyses

We used a logistic regression framework to link H5N8 presence and absence to the predictors. As our data had low prevalence values (< 10%), we applied a bootstrapping procedure with 1,000 repetitions, selecting all HPAI H5N8 presence sites with an equivalent number of absence sites, to minimize the problem of data imbalance. The absence sites were randomly selected under the condition that each absence site was more than 20 km away from others, in order to avoid duplicated absences.

We first performed univariate analyses for both the 2016/17 and 2020/21 epidemics to explore the risk factors for H5N8 occurrence in wild birds or in poultry. We then compared these results to the 2005/06 H5N1 epidemic by re-analyzing the data extracted from a previous study (Huang et al. 2019b). We calculated the *p*-values based on the mean Z-value from 1,000 repetitions (Gilbert et al. 2008). The variables with a *p*-value of less than 0.05 were identified as risk factors, and were used as candidate variables to construct multiple regression models after assessing multicollinearity of variables. We assessed multicollinearity diagnostics based on Pearson correlation coefficients (r). For those highly correlated variables (r > 0.7), such as monthly temperature and precipitation variables, we selected the variable with the smallest p value in univariate analyses. Similarly, we selected the variable with smallest *p* value for the community variables calculated for both the whole community and the assemblages with only high-risk species. We here included human population density as the control factor in both univariate and further multiple regression analyses. Road density was not used as the control factor, as it was highly correlated with the mean human population density (r = 0.78) and showed higher p-values than population density in univariate analyses.

For H5N8 occurrence in wild birds and poultry in each epidemic, we constructed two partial multiple models: community partial model including only waterbird community variables, and agro-climatic partial model including only agro-climatic variables, as well as a global model including all variables in partial models. Using these models, we conducted variance partitioning analyses, based on models’ mean Nagelkerke pseudo-R^2^ (other pseudo-R^2^ measures did not qualitatively change the results), to determine the relative importance of community predictors and ago-climatic predictors in explaining the variation in H5N8 occurrence.

Finally, we fitted the final models for the 2016/17 epidemic, and tested their predictive powers for the 2020/21 epidemic by using 2016/17 data as the training dataset and 2020/21 data as the test dataset. The final models were constructed by conducting model selection based on the abovementioned global models, using a statistical regularization approach (i.e., LASSO) (Tredennick et al. 2021), where we calculated the importance of each predictor, measured as the percentage of 1000 repetitions in which its coefficient did not shrink to exactly zero due to penalty. In LASSO analyses, we applied a 10-fold cross validation (CV) to generate the optimal value of the regularization parameter (λ), and did not penalize the control factor (mean human population density). We then conducted backward selection by removing the non-significant variable with the least importance in LASSO. In addition, we also conducted traditional backward selection by removing the variable with the highest *p*-value in each step. For both H5N8 in wild birds and in poultry, these two approaches generated a similar final model. We reported the mean values of the area under the Receiver Operating Characteristic curve (AUC) for both training and test datasets as the indicator of model performance. Moreover, we tested for, using Moran’s *I* index, spatial autocorrelation of the residuals of the final models, and found little evidence of spatial autocorrelation (*Supplementary file 1h*). All statistical analyses were conducted in R 4.0.2.

## Data availability

H5N8 outbreak data: FAO (http://empres-i.fao.org/eipws3g/).

Waterbird census data: IWC Online database (http://iwc.wetlands.org).

Human density data: http://sedac.ciesin.columbia.edu/data/collection/gpw-v4.

Road density data: http://www.globio.info/download-grip-dataset.

Poultry density data: http://www.fao.org/ag/againfo/resources/documents/multimedia/popup/glw.html.

CRU Climate data: crudata.uea.ac.uk/cru/data/hrg.

Waterbody data: www.worldwildlife.org/pages/global-lakes-and-wetlands-database.

ESA land cover: http://www.esa-landcover-cci.org/?q=node/197.

The code supporting the results are available on Source Code File.

## Acknowledgements

We are thankful for the contributions of all national coordinators (https://www.wetlands.org/our-network/iwc-coordinators) and for the thousands of volunteers who participated in the annual International Waterbird Census. We acknowledge the various sources of national and international funding that contribute to the continuation of the IWC.

## Funding

This work was supported by the National Natural Science Foundation of China (31870400, 82073616), and the Priority Academic Programme Development (PAPD) of Jiangsu Higher Education Institutions.

## Competing interest

The authors declare no competing interests.

## Additional files

Supplementary file 1. Supplementary Tables. (a) Results of univariate analyses (b coefficient, and t-value) for HPAI occurrence in wild birds in the three epidemics. (b) Results of multiple regression models for H5N8 occurrence in wild birds in the 2016/17 epidemic under different thresholds of identifying absences. (c) Results of multiple regression models for H5N8 occurrence in poultry in the 2016/17 epidemic under different thresholds of identifying absences. (d) Results of the final model (logistic regression model) for H5N8 outbreak risk in wild birds and poultry in the 2016/17 epidemic using presences without poor match. (e) Countries (with the number of bird survey sites) from the International Waterbird Census included in our study. (f) Description and abbreviation of the predictors used in the analysis. Variables with asterisks were log-transformed in the analyses. (g) 137 waterbird species (with order, family and genus they belong to) used in our study. The nomenclature follow the IUCN Red List. The higher-risk species and bridge species are, respectively, highlighted in bold and in red. (h) Mean Moran I values of the residuals for the final models for H5N8 occurrence in wild birds and poultry in the 2016/17 epidemic using different thresholds for identifying pseudo absences.

## Figures Legends

**Figure 2—source data 1:** The summary results (including the mean regression coefficient *b* and standard error *se, Z*-values and corresponding *p*-values) for univariate analyses.

## Notes

### Competing Interest Statement

The authors have declared no competing interest.

